# The site-frequency spectrum associated with Ξ-coalescents

**DOI:** 10.1101/025684

**Authors:** Jochen Blath, Mathias Christensen Cronjäger, Bjarki Eldon, Matthias Hammer

**Author notes:** Corresponding author: Bjarki Eldon, Museum für Naturkunde, Leibniz Institut für Evolutions- und Biodiversitätsforschung, Invalidenstraße 43, 10115 Berlin, Germany.

## Abstract

We give recursions for the expected site-frequency spectrum associated with so-called *Xi-coalescents*, that is exchangeable coalescents which admit *simultaneous multiple mergers* of ancestral lineages. Xi-coalescents arise, for example, in association with population models of skewed offspring distributions with diploidy, recurrent advantageous mutations, or strong bottlenecks. In contrast, the simpler *Lambda-coalescents* admit multiple mergers of lineages, but at most one such merger each time. Xi-coalescents, as well as Lambda-coalescents, can predict an excess of singletons, compared to the Kingman coalescent. We compare estimates of coalescent parameters when Xi-coalescents are applied to data generated by Lambda-coalescents, and vice versa. In general, Xi-coalescents predict fewer singletons than corresponding Lambda-coalescents, but a higher count of mutations of size larger than singletons. We fit examples of Xi-coalescents to unfolded site-frequency spectra obtained for autosomal loci of the diploid Atlantic cod, and obtain different coalescent parameter estimates than obtained with corresponding Lambda-coalescents. Our results provide new inference tools, and suggest that for autosomal population genetic data from diploid or polyploid highly fecund populations who may have skewed offspring distributions, one should not apply Lambda-coalescents, but Xi-coalescents.

## Introduction

The coalescent approach, ie. the idea of considering the (random) ancestral relations of alleles sampled from natural populations, has provided rich mathematical theory [cf. 7], and very useful inference methods (cf. eg. [18, 45] for reviews). Initiated by the Kingman coalescent [28, 30, 29], coalescent models now include the family of Lambda-(Λ-)coalescents [35, 36, 17], and Xi-(Ξ-)coalescents [40, 33, 37]. Ξ-coalescents admit *simultaneous multiple mergers* of ancestral lineages. Thus, in each merger event, distinct groups of ancestral lineages can merge at the same time, and each group can have *more* than two lineages. Λ-coalescents, in contrast, only allow one group - possibly containing more than two lineages - to merge each time. Thus, due to multiple mergers, the derivation and application of inference methods becomes harder as one moves from the Kingman coalescent to Lambda-coalescent models, and from Λ-coalescents to Ξ-coalescents. Ξ-coalescents can be obtained from diploid population models [9, 34, 14]. They also arise in models of repeated strong bottlenecks [11], and in models of selective sweeps [19, 20].

[24] obtained closed-form expressions for the expected site-frequency spectrum, and (co)-variances, when associated with the Kingman coalescent. [10] obtain recursions for expected values and (co)-variances when associated with Λ-coalescents. However, the complexity of the recursions means that (co)-variances, when associated with Λ-coalescents, can only be computed for small sample sizes. The expected values can be applied in distance statistics [25], and in an approximate likelihood approach [10, 22].

Multiple merger coalescents can be obtained from population models that admit high fecundity and skewed offspring distributions, characteristics associated with many marine populations [1, 23, 8, 12, 13, 26, 27, 38]. Indeed, [2] find much better fit (ie. minimizing the sum of the squared distance between observed and expected values) between data on autosomal genes from Atlantic cod and Λ-coalescents than with the Kingman coalescent. Simulation results of [38] suggest that the site-frequency spectrum of (at least some) Ξ-coalescents is multi-modal, a pattern observed in data on the autosomal *Ckma* gene in Atlantic cod [2]. Based on this evidence, a way to compute expected values of the site-frequency spectrum associated with Ξ-coalescents should be a welcome and important addition to the set of inference methods for population genetics. Time-changed Kingman-coalescents obtained from models of population growth can predict an excess of singletons, a characteristic observed for some marine populations [cf. eg. 27]. However, the right tails of the site-frequency spectrum predicted by multiple-merger coalescents and time-changed Kingman-coalescents differ, which makes it possible to distinguish between them [22].

In this work, we obtain recursions for the expected site-frequency spectrum associated with general Ξ-coalescents, with an approach similar to the one applied by [10]. We compare estimates of coalescent parameters when applied to simulated data obtained under Λ-coalescents, and vice-versa. Since the recursions for the expected values are already fairly complex, and computationally expensive, we expect recursions for the (co)-variances to be even more so. The (co)-variances will therefore not be addressed.

We estimate coalescent parameters associated with specific examples of Xi-coalescents for the unfolded site-frequency spectrum of 3 autosomal loci [2] of the highly fecund Atlantic cod. Our simple method involves minimising the distance between observed and expected values, where the distance is not calibrated by the corresponding variance. Our estimates differ from previous estimates obtained with the use of Lambda-coalescents. The main biological implication of our results are that Xi-coalescents should be applied to autosomal data from highly fecund diploid (or polyploid) populations, and Lambda-coalescents to haploid data such as mitochondrial DNA.

The paper is structured as follows: First we give a precise mathematical description of the various coalescent models. We then state our main result on the expected site frequency spectrum of Ξ-coalescents, Theorem [2]. This is followed by a discussion of some specific examples of Ξ-coalescents. Some numerical examples which illustrate the difference in the site-frequency spectrum between Lambda- and Xi-coalescents are then presented, followed by an application to autosomal Atlantic cod data. The proofs are collected in an Appendix.

## Theory

For ease of reference, we collect the notation we use in the following table.

**Table 1:**
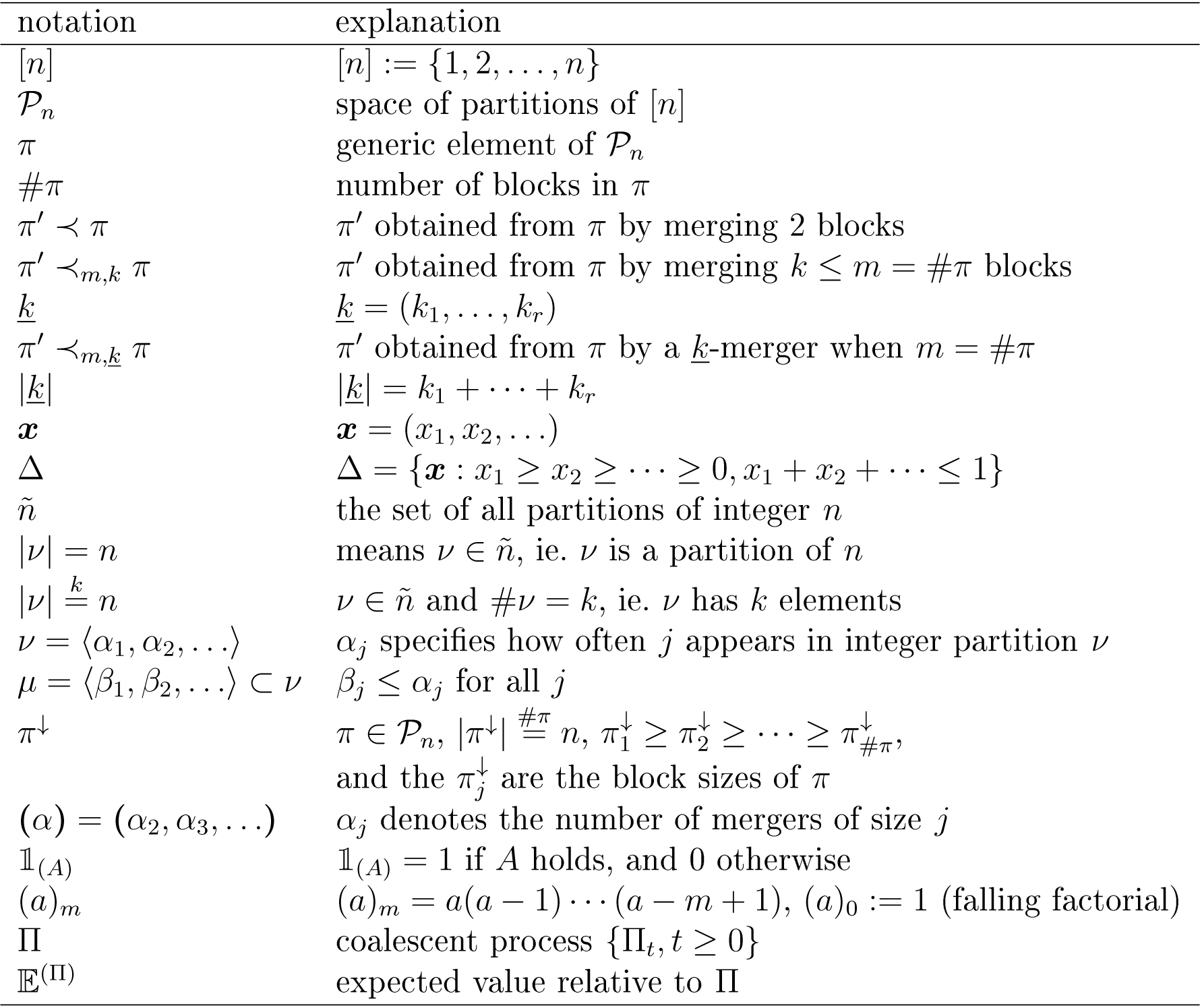
Notation.

### Coalescent models

We briefly review the basic coalescent models, namely the Kingman-, Λ-, and Ξ-coalescents. They all have in common to be continuous-time Markov chains, taking values in the space of partitions of the natural numbers 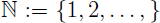, whose restriction to the first *n* integers can be described as follows: Let *𝒫_n_* denote the space of partitions of 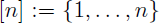. We write π for a generic element of *𝒫_n_*, and *#π* for the *size* of *π*, i.e. for the number of blocks 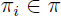. Thus for 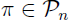 we have 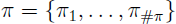 with 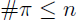.

### Kingman coalescent

If *π*, 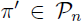 with 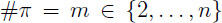, we write 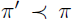 if there exist *i*, 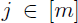 with 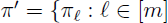, 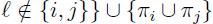, ie. π^′^ is obtained from *π* by merging blocks and *π_i_* and *π_j_*. If the transition rates of the continuous-time Markov chain 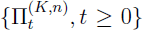 with values in *𝒫_n_*, and starting from state 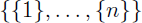 at time *t* = 0, are given by

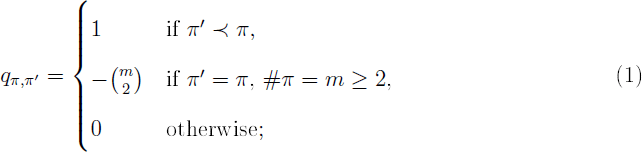

we refer to 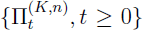 as the Kingman-*n*-coalescent. The process is stopped at time inf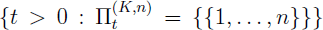, ie. when the most recent common ancestor of the *n* lineages has been reached.

### Lambda-coalescent

If *π*, 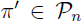 with 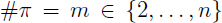, and there exist indices 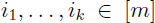 with 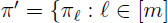, 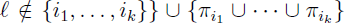, we write 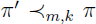 and say that a *k*-merger has occurred, with 2 ≤ *k* ≤ *m*. For a finite measure Λ on [0, 1], define

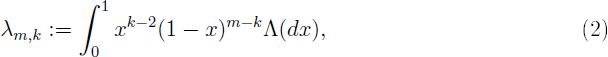

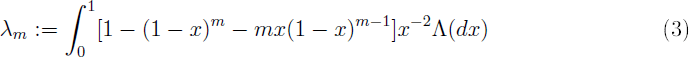

if the integral in (3) exists, and

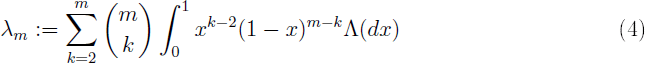

otherwise. For example, the integral in (3) does not exist in ease of the Beta (2 – α, α)-coaleseent introduced by [41].

A *𝒫_n_*-valued continuous-time Markov chain 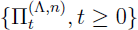 with transition rates *q_π,π′_* from π to π′ given by

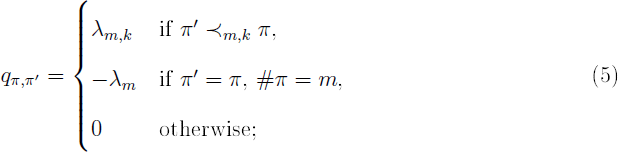

is referred to as a Λ-*n*-coalescent. The waiting time in state π is exponential with rate λ*_m_* as in (3, 4).

### Xi-coalescent

Now we specify the transition rates for a Ξ-*n*-coalescent. Let 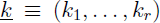 denote a vector of positive integers of length *r* ≥ 1. We write 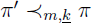 if 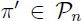 with 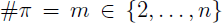, and there exist 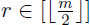 groups of indices 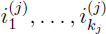, *j* = 1, …, *r*, such that

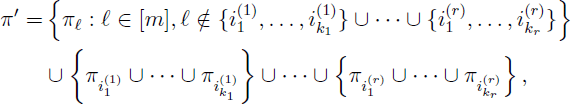

by which we denote a transition where blocks with indices 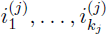 merge into a single block, for 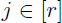. Thus, a transition denoted by 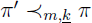 is a *simultaneous multiple merger*, where 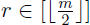 such mergers occur simultaneously. The vector 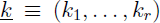 specifies the merger sizes, and we write 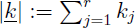.

Let Δ denote the infinite simplex

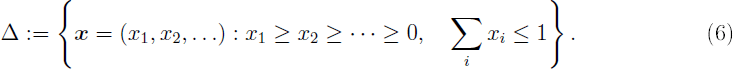

Let 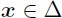, 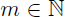, 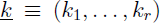 with *k_i_* ≥ 2 and 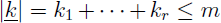 the sum of the *r* merger sizes, and 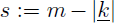 the number of blocks unaffected by the merger(s) specified by *<u>k</u>*. Define the functions *f* (*x*, *m*, *<u>k</u>*) and *g*(*x*, *m*, *<u>k</u>*), with 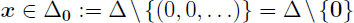,

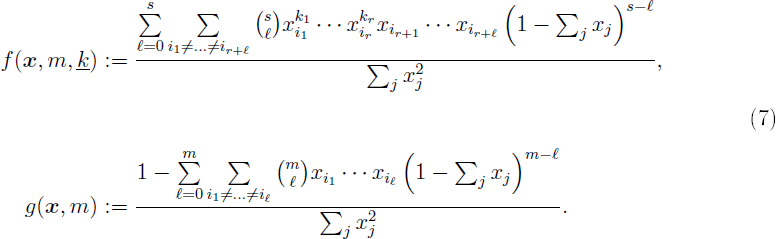

Let Ξ_0_ denote a finite measure on Δ_0_, and write 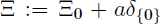. Further, let 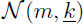 denote the number of ways of arranging *m* items into *r* non-empty groups whose sizes are given by *<u>k</u>*. With ℓ*_j_* denoting the number of *k*_1_, …, *k_r_* equal to *j*, one checks that, with 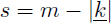 [40],

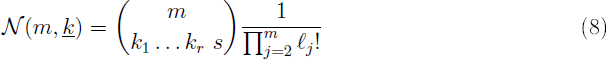

(recall that 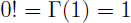). Now define [40], with 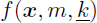 and *g*(*x*,*m*) given by (7),

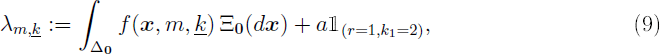

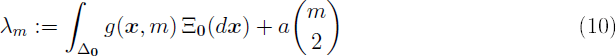

if the integral in (10) exists, and

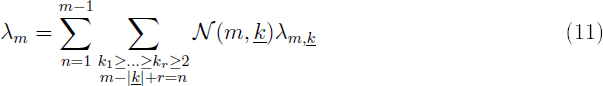

otherwise. A continuous-time *𝒫_n_*-valued Markov chain with transitions *q_π,π'_* given by [40]

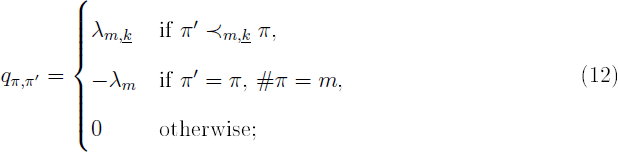

is referred to as a Ξ-*n*-coalescent, and denoted by 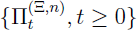. The waiting time in state π is exponential with rate λ*_m_* as in (10, 11).

### Specific examples of Xi-coalescents

Out of the rich class of Xi-coalescents, several special cases have been identified either for biological / modeling relevance or mathematical tractability. The example most relevant for us is concerned with diploidy.

Haploid population models are probably the most common models in mathematical population genetics. Diploidy, and other forms of polyploidy, are, however, widely found in nature. Atlantic cod is diploid, and oysters show both tetraploidy and triploidy [cf. eg. 32]. In polyploid models which admit skewed offspring distribution, one should observe up to *M* ≥ 2 simultaneous mergers, where *M* is some fixed number which reflects the level of polyploidy. Indeed, [34] model diploidy in a population with a general skewed offspring distribution, and obtain a Xi-coalescent, which admits simultaneous multiple mergers in up to four groups.

Indeed, a mathematical description of a Xi-coalescent which admits up to *M* simultaneous mergers is as follows. Take a finite measure Λ on [0, 1] (which would normally describe a Λ-coalescent). For convenience, let 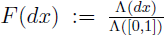 be the corresponding normalized probability measure. Then with *M* ≥ 2, define the measure Ξ on the simplex Δ by

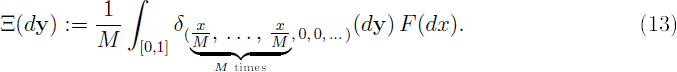

The interpretation is this: If the normalized Lambda-measure *F* produces a multiple merger event, in which individual active ancestral lineages (blocks of the current partition) take part with probability 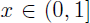, then the participating lineages are randomly grouped into *M* simultaneous mergers (each with probability 1/*M*). Observe that if 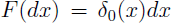, then (13) becomes 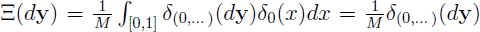, which corresponds to a Kingman-coalescent with time scaled by a factor 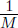.

For a given merger of the Ξ-coalescent into 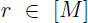 groups of sizes given by 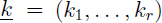, with 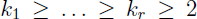 and 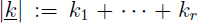, when *m* active ancestral lineages are present, with 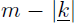 lineages unaffected by the given merger, the transition rates are given by

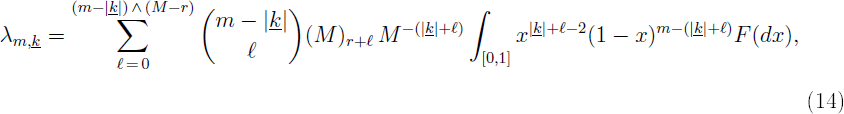

where 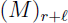 denotes the falling factorial. The rates (14) depend on the choice of the probability measure *F* determined by the underlying population model. A proof of (14) is in the Appendix. We say that a Xi-coalescent is *M-fold* if the transition rates are given by (14).

Xi-coalescents which admit at most *M* = 4 simultaneous mergers arise from diploid Cannings population models with skewed offspring distribution as shown in [34, 9], and are thus relevant for population genetics. In fact, if a haploid model (for example for mitochondrial DNA) is governed by a Λ-coalescent, the corresponding diploid model (concerning the core genome) might naturally lead to Xi-coalescents. Indeed, [9] derive a 4-fold Xi-coalescent from a diploid model, in which exactly one pair of diploid parents contribute diploid offspring in each reproduction event. Hence, since 4 parental chromosomes are involved in each event, one can observe up to 4 simultaneous mergers. This was also observed by [34] in association with a diploid population model, but under a more general reproduction law than considered by [9]. Xi-coalescents are also classified by [14] for a very general diploid exchangeable Cannings model in which arbitrary pairs of diploid parents contribute offspring in each reproduction event. This generalises the model by [34], in which each individual forms at most one parental pair in each generation. For a detailed classification of coalescent limits, see [34, 37, 33, 14].

In truly diploid models, as considered by [34, 9], selling is excluded, which leads to a ‘separation of timescales’ phenomenon in the ancestral process, in which blocks which reside in the same diploid individual instantaneously ‘disperse’; thus the configuration of blocks in diploid individuals becomes irrelevant in the ancestral process (see Cor. 4.3 in [34]).

A natural candidate for *F* may be the beta distribution with parameters ν > 0 and γ > 0 (cf. e.g. [9]), with density

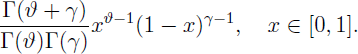

In this case, the rate 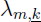 in (14) takes the form

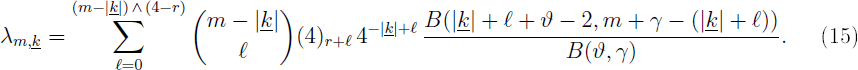

A different choice is based on a model of Eldon and Wakeley [23], where

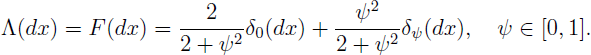

In this case, the rates reduce to

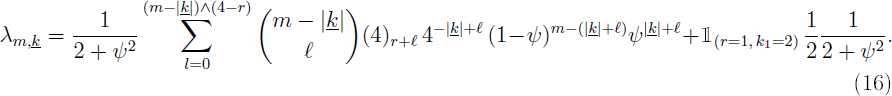

In (16) the parameter ψ has a clear biological interpretation as the fraction of the diploid population replaced by the offspring of the reproducing parental pair in one generation. The interpretation of the parameters in (15) is perhaps less clear.

A multi-loci ancestral recombination graph in which simultaneous mergers are admitted is obtained by [9] in which the framework of [34] is borrowed. There, one can think of the reproduction model as a two-atom Lambda-measure, one atom at zero, and another at some point 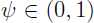. If the atom at ψ has mass of order at most *N*^–2^, the limit process admits simultaneous mergers. The order *N*^–2^ represents the order of the expected time which two gene copies need to coalesce when only 1 diploid offspring is produced in each reproduction event. In [9], complete dispersion of chromosomes also occurs, and the configuration of blocks among diploid individuals becomes irrelevant in the limit process.

Xi-coalescents can also be obtained from a population model where the population size varies substantially due to recurrent bottlenecks. This has been introduced and discussed in [11], who obtain a randomly time-changed Kingman coalescent, which thus yields a Xi-coalescent.

Durrett and Schweinsberg [19, 20] show that a Xi-coalescent gives a good approximation [20, cf. Prop. 3.1] to the genealogy of a locus subject to recurrent beneficial mutations.

These examples suggest that Xi-coalescents form an important class of mathematical objects with which to study genetic diversity.

### The site-frequency spectrum

The site-frequency spectrum is a simple summary statistic of the full DNA sequence data, but contains valuable information about variation among individuals. We assume the infinitely-many sites mutation model [46], in which mutations occur as independent Poisson processes on the branches of a given gene genealogy with rate θ (or θ/2) for some constant θ > 0, and no two mutations occur at the same site. The constant θ is determined by the ratio μ/*c_N_*, where μ is the per-generation mutation rate, and *c_N_* is the probability of two distinct individuals (gene copies) sharing a common ancestor in the previous generation. We refer to [22] for a discussion of the relation between mutation and timescales of different coalescent processes.

Given sample size *n*, we let 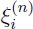 denote the number of polymorphic sites at which one variant (the derived mutation) is observed in *i* copies. The random vector

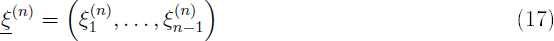

is known as the (unfolded) *site-frequency spectrum*. If information about ancestral states is unavailable, so that one does not know which variant is new, one considers the *folded* spectrum spectrum 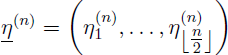 in which

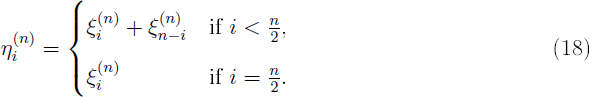

[24] obtains closed-form solutions for expected values and (co)-variances of the site-frequency spectrum associated with the Kingman coalescent П^(*K*)^. Indeed [24],

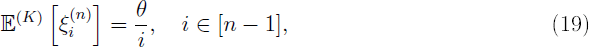

where 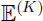 denotes expectation with respect to Kingman coalescent.

Let 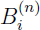 denote the random total length of branches subtending 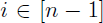 leaves. By leaves we refer to the special kind of vertices in the random graph generated by a given coalescent process which would represent the sampled DNA sequences after mutations have been added to the graph. Since the mutation process can be separated from the genealogy (the random graph) it can sometimes be useful to consider properties of the graph itself. Result (19) follows from [24],

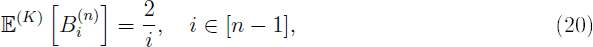

and the infinitely-many sites mutation model.

Due to the multiple merger property of Λ- and Ξ-coalescents, closed-form expressions for 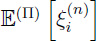 for 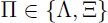 are quite hard to obtain. A key quantity in computing 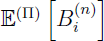 for multiple merger coalescents is

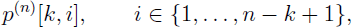

which can be described as the probability that starting from *n* blocks, conditioned that there are at some point in time exactly *k* blocks, one of them, sampled uniformly at random, subtends 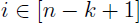 leaves. See Figure 1 for an illustration.

**Figure 1:**
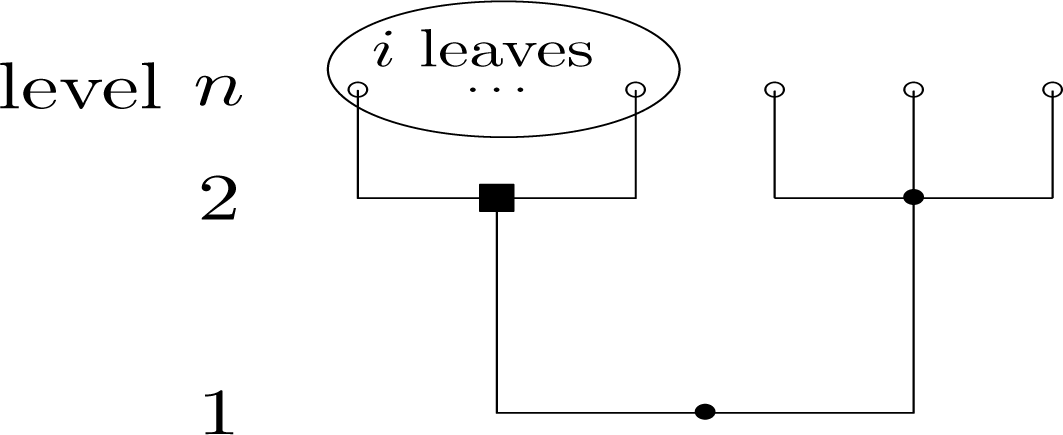
An illustration of an event with probability 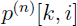, with *k* = 2, and 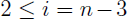 for *n* leaves (shown as open circles). In the example shown, the process reaches *k* = 2 blocks in one transition, which is simultaneous merger involving all the *i* leaves (encircled) as shown. All the *i* leaves are subtended by the block represented by a black square. A ‘level’ refers to the values taken by the block represented by a black square. A ‘level’ refers to the values taken by the block-counting process; filled symbols represent ancestral blocks.

With *g*(*n, k*) we denote the expected length of time during which we see 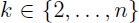 blocks, given that we started from *n* ≥ 2 blocks. Given 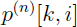 and *g*(*n, k*), 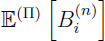 can be computed as follows:

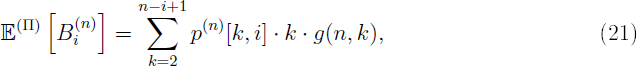

where moreover *g*(*n, k*) can be computed recursively. This is shown in [10] for Λ-coalescents but in fact holds for Ξ-coalescents as well. Hence, it suffices to obtain a recursion for 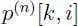. For the Λ-case, [10] obtain the recursion

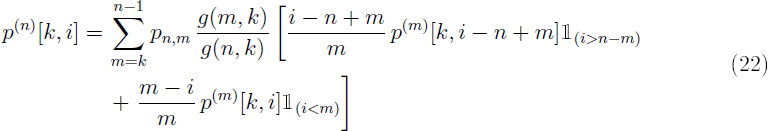

in which *p_n, m_* is the probability that the block-counting process jumps from *n* to *m* blocks.

Before we turn to the expected site-frequency spectrum associated with Ξ-coalescents, we give the recursion to compute *g*(*n, k*).

**Lemma 1.** *Let p_n,m_ denote the probability that the block-counting process associated with a Ξ-n-coalescent with transition rates* (12) *jumps from n* ≥ 2 *to* 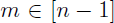 *blocks. For any* 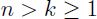, *we have*

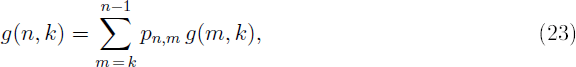

*with the boundary condition*

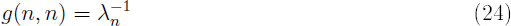

*for any n* ≥ 2, *where* 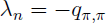 *with* 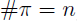, *see (10, 11, 12).*

A proof of Lemma 1 is given in the Appendix.

### The expected site-frequency spectrum associated with Ξ-coalescents

Since formula (21) holds for any exchangeable coalescent, it suffices to obtain a recursion for 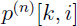 when associated with Ξ-coalescents. Before we state the main result, we review our notation for partitions of positive integers. Partitions of integers are helpful in enumerating the different ways in which the number of active blocks can change in one transition in a Xi-coalescent.

A *partition* of 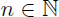 is a non-increasing sequence of positive integers whose sum is *n*. By ñ we denote the set of all partitions of *n*, we denote by ν a generic element of *ñ*. By way of example,

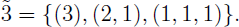

If 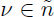, we write 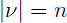. The *size* of a partition ν is defined as the length of the sequence, and is denoted by #ν. If 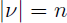 with size 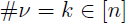, we write 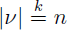. Thus, 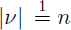 if and only if ν = (*n*); 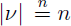 if and only if 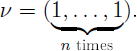.

Another way of representing an integer partition ν is by specifying how often positive integer 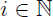 appears in ν (see [15] for details). Thus, we will also denote ν by 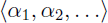, where *α_i_* denotes the number of times integer *i* appears in the given partition ν. A partition 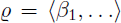 is a *sub-partition* of 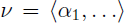, denoted 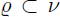, if and only if 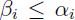 for all *i*. For a set partition 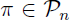, we define the *integer partition associated with* π, denoted 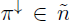 as the partition of *n* obtained by listing the block sizes of π in decreasing order. More detailed discussion of partitions of integers can be found eg. in [44].

The role of integer partitions in association with Ξ-coalescents should now be clear. We can enumerate all the possible ways the block counting process can jump from *n* to *m* active blocks by specifying the partitions of *n* (see eg. [15] for details). The elements of the sequence 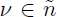 specify the merger sizes, with the obvious exclusion of mergers of size 1. By way of example, integer partition 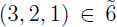 specifies a simultaneous merger of 3 blocks and 2 blocks, and one block remains unchanged, when we have 6 active blocks. By (12), any such transition happens at rate λ_6,(3,2)_.

More generally, given integer partition 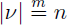 with 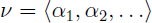, put 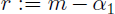 for the number of elements of the sequence that are larger than 1, so that

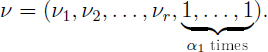

Then 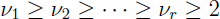 2, and defining 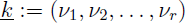, the corresponding transitions in which 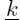 specifies the sizes of the *r* mergers involved happen at rate 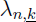, see (12). Moreover, the probability 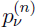 of such a transition is given by [15, Lemma 2.2.2]

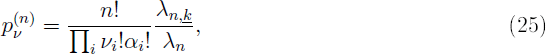

where the combinatorial factor

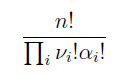

(coinciding with 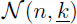 as introduced in (8)) denotes the number of different ways of merging *n* blocks in *r* groups specified by 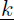, and 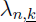, λ*_n_* are given by (9, 10, 11).

Now we state our main theorem, which contains the recursion for 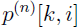 needed to compute 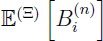. The theorem holds for all Ξ-*n*-coalescents whose block-counting process visits every possible state with positive probability, which is true of all examples of Xi-coalescents that we consider. The assumption is not very restrictive, since it excludes only degenerate cases where the measure Ξ is concentrated on a vertex of the infinite simplex Δ_0_, see Proposition 6 in the Appendix for a precise formulation.

**Theorem 2.** *[15] Let* 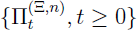 *be a Ξ-n-coalescent with transition rates* (12) *such that the corresponding block-counting process hits every k* ∈ [*n*] *with positive probability. Then, for* 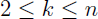 *and* 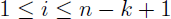, *we have*

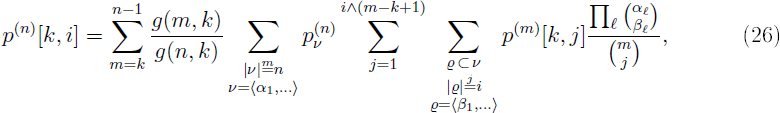

*with the boundary cases* 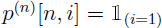.

A proof is provided in the Appendix, along with a necessary and sufficient condition for the block-counting process to hit state 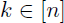 with positive probability (see Proposition 6).

The recursion (26) for 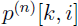 simplifies when one restricts to Ξ-coalescents with at most *M* ≥ 2 simultaneous mergers (*M* = 1 simply gives a Lambda-coalescent), as shown in Cor. 3. Before we state Cor. 3, we briefly review *ordered mergers*. The computations are more efficient for Xi-coalescents restricted to at most *M* simulataneous mergers, and when one considers ordered mergers rather than partitions of integers. Let

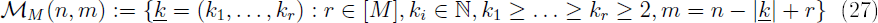

denote the set of single and up to *M* simultaneous ordered mergers by which the block-counting process can jump from *n* ≥ 2 to 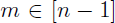 blocks in 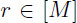 mergers. The set 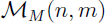 corresponds to the set of all integer partitions 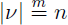 such that, if 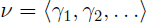, we have 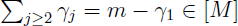. Indeed, for 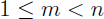, we have a bijection

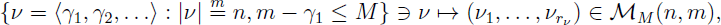

with 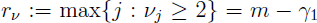, for 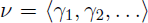. Obviously, the inverse bijection is given by

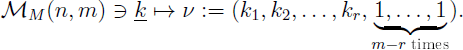

Informally, ordered mergers are just integer partitions where all elements equal to one are omitted.

For ease of presentation, we will also write 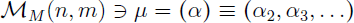 where *α_j_* denotes the number of occurrences of mergers of size *j* in *μ*. By 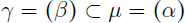 we denote a *submerger* γ of μ where 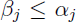 for all *j, including* the case *β_j_* = 0. Finally, the *size* of 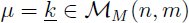 is just the length of the sequence 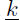, i.e. 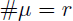 for 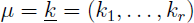.

If μ is the ordered merger corresponding to some integer partition 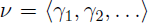, 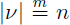 as above, then 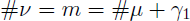.

**Corollary 3.** *Let* 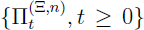 *be a Ξ-n-coalescent with transition rates* (12) *such that at most M* ≥ 2 *simultaneous mergers are possible, and such that the corresponding block-counting process hits every* 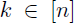 *with positive probability. Then, for* 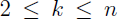 *and* 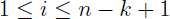, *we have*

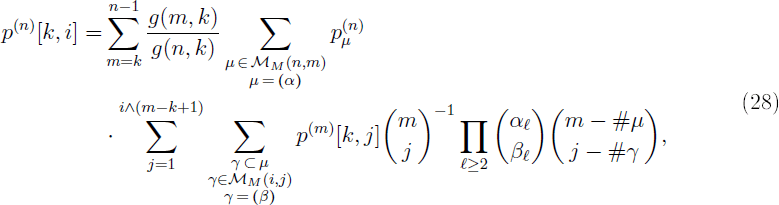

*with the boundary cases* 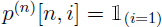.

Informally, the first sum in recursion (28) is over the number of blocks the block-counting process can jump to, given that it starts in *n*, and conditioned on it hits *k*. The second sum is over all (up to *M* simultaneous) mergers (μ) in which one can jump from *n* to *m* blocks, and the last sum is over all the ways mergers involving the *i* leaves can be nested within each given merger μ.

Corollary 3 is simply another way of representing 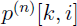 (26) (see Thm, 2) in terms of ordered mergers. The switch in focus to ordered mergers from integer partitions obliviates the need to keep track of all the 1s. Also, since we restrict to Ξ-coalescents which admit at most *M* simultaneous mergers, the required mergers can be generated quite efficiently.

The computation of 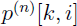 can be checked by noting that for each fixed *n* ≥ 2, and with 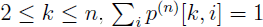. One can also compute 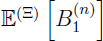 with a simple recursion as follows. Let 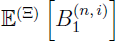 denote the expected length of *i* external branches, given *n* ≥ 2 active blocks; ie. When we start from *n* active blocks, *i* of which are singleton blocks. Define, for all 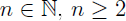,

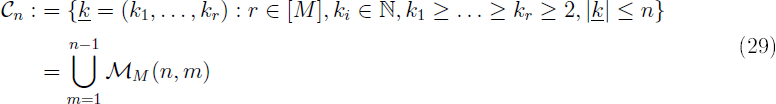

as the set of all ordered (up to *M* simultaneous) mergers given *n* active blocks, where 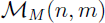 was defined in (27).

**Lemma 4.** *Let* 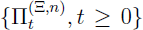 *be a* Ξ*-n-coalescent with transition rates* (12) *such that at most M simultaneous mergers are possible. With* λ*_n_* given by (10, 11), let C_n_ be given by (29), *and let* 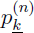 denote the probability of merger 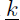 *when the number of active blocks is n. Then*,

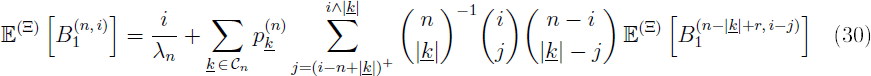

*with the boundary condition* 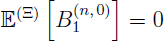.

## Numerical results

### Expected branch lengths 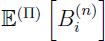

Under the infinitely many sites mutation model, the expected site-frequency spectrum is given by 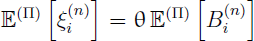 were θ < 0 is the appropriately scaled mutation rate. Hence, it suffices to consider 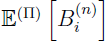 in a comparison of the site-frequency spectrum associated with different coalescent models. In Figure 2, we consider the 4-fold Ξ-coalescent, when the measure *F* in (14) is associated with the beta-density, with 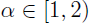 [41],

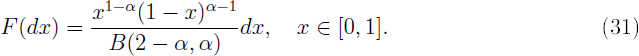

**Figure 2:**
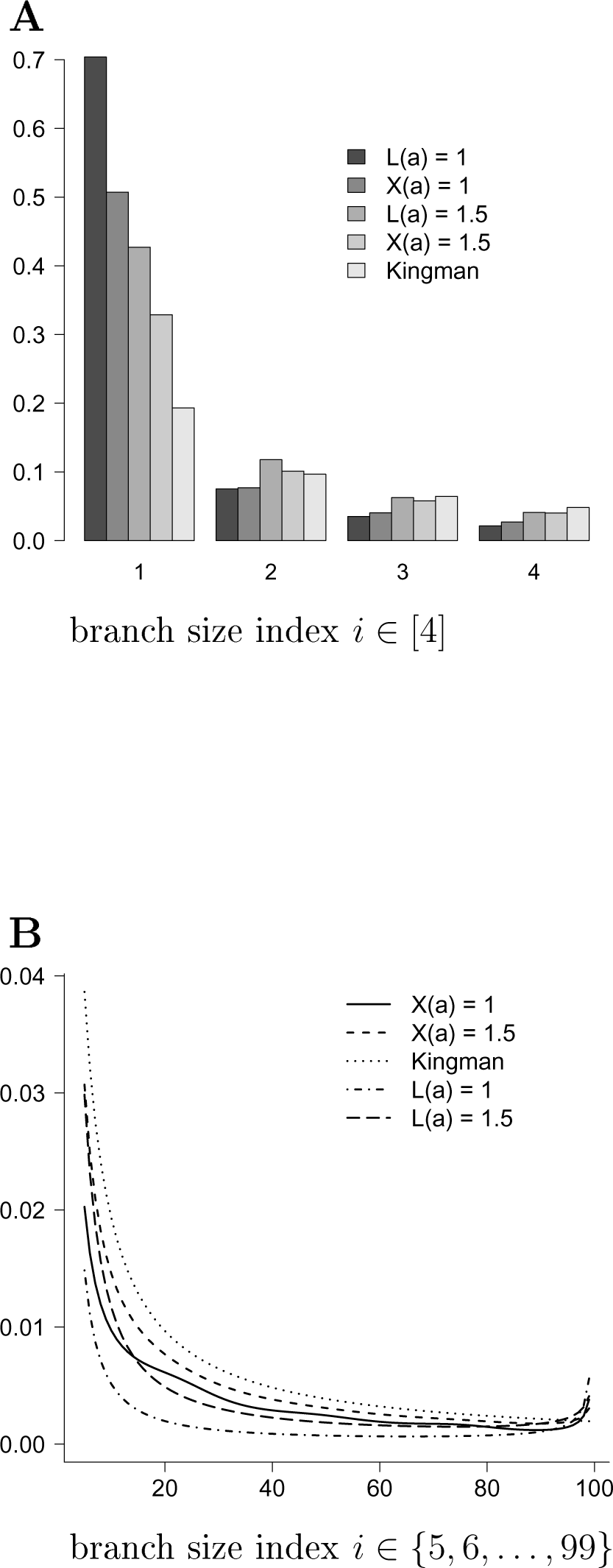
The relative expected lengths 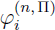 (32) for *n* = 100 and coalescent process П as follows: `L(a)` denotes the Lambda-Beta coalescent with parameter value a (α) as shown, and `X(a)` denotes the Xi-Beta coalescent with parameter value a (α) as shown. In **A**, only the first 4 classes are shown; in **B**, the remaining classes.

The range of α is the interval [1. 2), since for 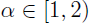, one obtains a Lambda-Beta coalescent from a supercritical branching population model [41]. We do not have a microscopic diploid population model which explicitly yields a Xi-Beta coalescent. However, the results of [34] indicate that such a process should exist. Also, the Lambda-Beta coalescent is one of the most studied examples of Lambda-coalescents. However, the existence of the Xi-Dirac coalescent was proved by [9]. The expected branch lengths associated with a Lambda-coalescent with *F*- measure (31), as well as the Kingman coalescent, are also shown in Figure 2 for comparison. We consider the normalised expected spectrum

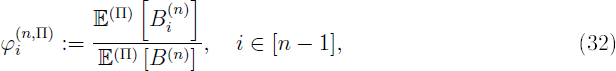

in which 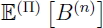 is the expected total size of the genealogy 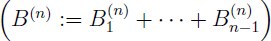. Define 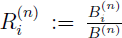. Let 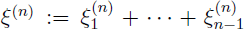 denote the (random) total number of segregating sites, and define the normalised spectrum 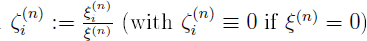. The reasons for our preference for 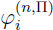 over 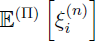 are the following. The quantity 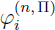, which is an approximation of the expected normalised spectrum 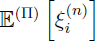 is, clearly, *not* a function of the mutation rate θ. The expected normalised spectrum 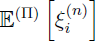 is well approximated by 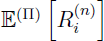, and is also quite robust to changes in mutation rate, if the mutation rate is not very small [22]. Since 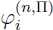 is a decent approximation of 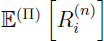 (results not shown), 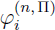 is therefore a good approximation of 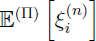. One can therefore use 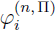 to estimate coalescent parameters, for example by minimising sum-of-squares, without the need to jointly estimate the mutation rate.

Refer to a Xi-coalescent with marginal (single-locus) rate described by Eq. (14) in [9] as the Xi-Dirac-Kingman coalescent. This corresponds to a process with coalescent rates given by (14), where the measure *F* has an atom at some 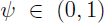, but there is an additional atom at 0, which represents the ‘Kingman component’. This process was derived by [9] in connection with ancestral recombination graphs involving multiple loci, in which simultaneous multiple mergers are admitted. In some cases, even for ψ close to 1, the associated 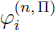 is very similar 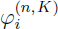 (results not shown).

In Figure 2 the 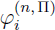 for corresponding Lambda-Beta and Xi-Beta coalescents are compared. As Figure 2 shows, the two processes predict different patterns of the site-frequency spectrum, at least for 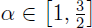. Both processes can predict a significant excess of singletons relative to the Kingman coalescent.

Similar conclusions can be reached from Figure 3 in which the measure *F* is given by the Dirac-measure 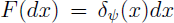 for some 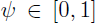 (no Kingman component). The Xi-Dirac-coalescent, for ψ = 0.95, displays a multimodal graph of 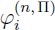, but the relative ‘height’ of the modes is small (< 1% of the total expected length). As well as an excess of singleton polymorphisms, [2] observe multi-modality, or small ‘bumps’, in the site-frequency spectrum associated with the *Ckma* gene in Atlantic cod.

**Figure 3:**
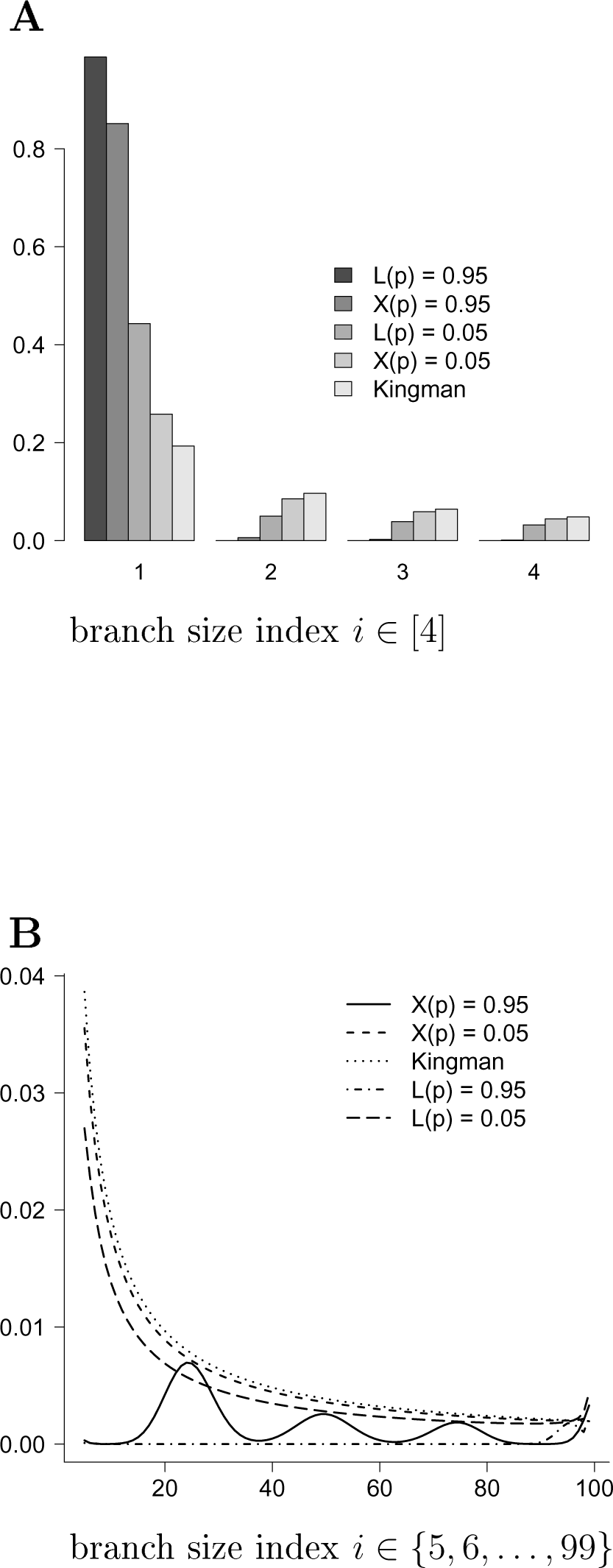
The relative expected lengths 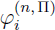 (32) for *n* = 100 and coalescent process П as follows: `L(p)` denotes the Lambda-Dirac coalescent with parameter value `p` (α) as shown, and `X(p)` denotes the Xi-Dirac coalescent with parameter value `p` (α) as shown. In **A**, only the first 4 classes are shown; in **B**, the remaining classes.

### Coalescent parameter estimates

As Figures 2 and 3 indicate, Xi- and Lambda-coalescents predict different site-frequency spectra. In Table 2 we record parameter estimates obtained when the ‘data’ are branch *B_i_* simulated under either a 4-fold Xi-coalescent (14) or a Lambda-coalescent (5) with parameter values as shown. However, we consider an extensive comparison of different examples of the large class of Xi-coalescent beyond the scope of the current work. Let ν denote a generic coalescent parameter. If 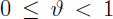, 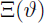 denotes a 4-fold Xi-Dirac coalescent, with *F*-measure 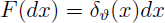 in (14), and 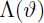 a Lambda-Dirac coalescent. If 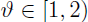, 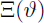 denotes a 4-fold Xi-Beta coalescent, and 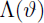 a Lambda-Beta coalescent. Estimates of ν attributed to coalescent process П_2_ are obtained with an ℓ_2_ norm applied to the normalised lengths 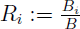 (where we drop the superscript ‘(n)’) drawn from coalescent process П_1_, and 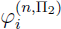,

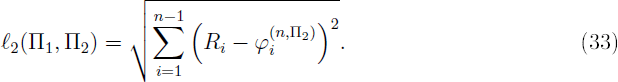

**Table 2:** Estimates of coalescent parameter ν, obtained by the use of the ℓ_2_ norm (33), when the ‘data’ (branch lengths of a realized genealogy) are obtained by a Lambda- 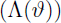 or a Xi-coalescent 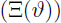 as shown. In the top half, the ‘data’ is generated by a 4-fold Xi-coalescent with parameter value as shown, and a parameter estimate obtained for the corresponding Lambda-coalescent. In the bottom half, branch lengths are generated from a Lambda-coalescent with parameter values as shown, and parameter estimates obtained for the corresponding 4-fold Xi-coalescent. Parameter estimates (mean 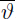; standard deviation 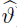) obtained for *n* = 50 leaves from 10^4^ replicates. Estimates were obtained over the grids 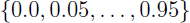 and 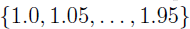.

| $\Pi(\vartheta)$ | $\bar{\vartheta}$ | $\widehat{\vartheta}$ |
| --- | --- | --- |
| $\Xi(0.05)$ | 0.02 | 0.029 |
| $\Xi(0.95)$ | 0.30 | 0.225 |
| $\Xi(1.0)$ | 1.44 | 0.257 |
| $\Xi(1.5)$ | 1.70 | 0.208 |
| $\Lambda(0.05)$ | 0.23 | 0.144 |
| $\Lambda(0.95)$ | 0.95 | 0.035 |
| $\Lambda(1.0)$ | 1.01 | 0.041 |
| $\Lambda(1.5)$ | 1.20 | 0.256 |

If the *B_i_* are draw from a Xi-Dirac coalescent (П_1_), we estimate ν associated with a Lambda-Dirac coalescent (П_2_), and vice versa. If the *B_i_* are drawn form a Xi-Beta coalescent (П_1_), we estimate ν associated with a Lambda-Beta coalescent (П_2_), and vice versa. The ℓ_2_ norm (33) is appealing since it makes no assumptions about the distribution of *R_i_*. In contrast, the Poisson Random Field (PRF) model [39] assumes Poisson distribution of mutation counts. Indeed, [16] prove that the law of the joint site-frequency spectrum converges, in the limit of infinite sample size, to that of independent Poissons, when associated with the Kingman coalescent. The asymptotic results of [4, 6, 5] show that multiple-merger coalescents belong to completely different asymptotic regimes than the PRF model assumes.

A more suitable distance statistic, similar to the *G_ξ_* statistic suggested by [25, Eq. (8)], could be

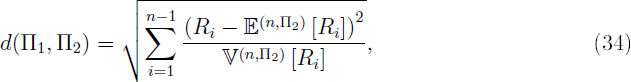

where 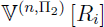 denotes the variance of *R_i_* computed with respect to П_2_. However, we can neither represent 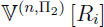 nor 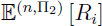 as simple functions of ν or *n*. In actual applications, one would replace *R_i_* in (34) with 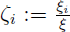 the normalised site-frequency spectrum (normalized by the total number of segregating sites ξ).

As Table 2 shows, a Lambda-Dirac coalescent underestimates ψ when the data are generated by a Xi-Dirac coalescent, and Lambda-Beta overestimates α when the data are generated by a Xi-Beta coalescent. When we switch the generation of data from Xi- to Lambda-coalescent, we reach the opposite conclusions. A Xi-Dirac coalescent overestimates the parameter (ψ) when the data are generated by a Lambda-Dirac coalescent, and the Xi-Beta coalescent underestimates α when the data are generated by a Lambda-Beta coalescent. The ℓ_2_ norm (33) recovers the true true parameter values reasonably well the model is correctly specified (Table 4).

The difference between the corresponding Xi- and Lambda-coalescent are further illustrated in Figure 4, in which the distance between the normalized expected spectra 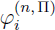 (32), with П as shown, is quantified by the ℓ_2_ norm (33). The graphs in Figure 4 show clearly that even when the parameters associated with the corresponding Xi- and Lambda-coalescents are the same, the difference in 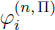 can be substantial (except of course when the Lambda-coalescent is the Kingman coalescent; which happens when α = 2 or ψ = 0).

**Figure 4:**
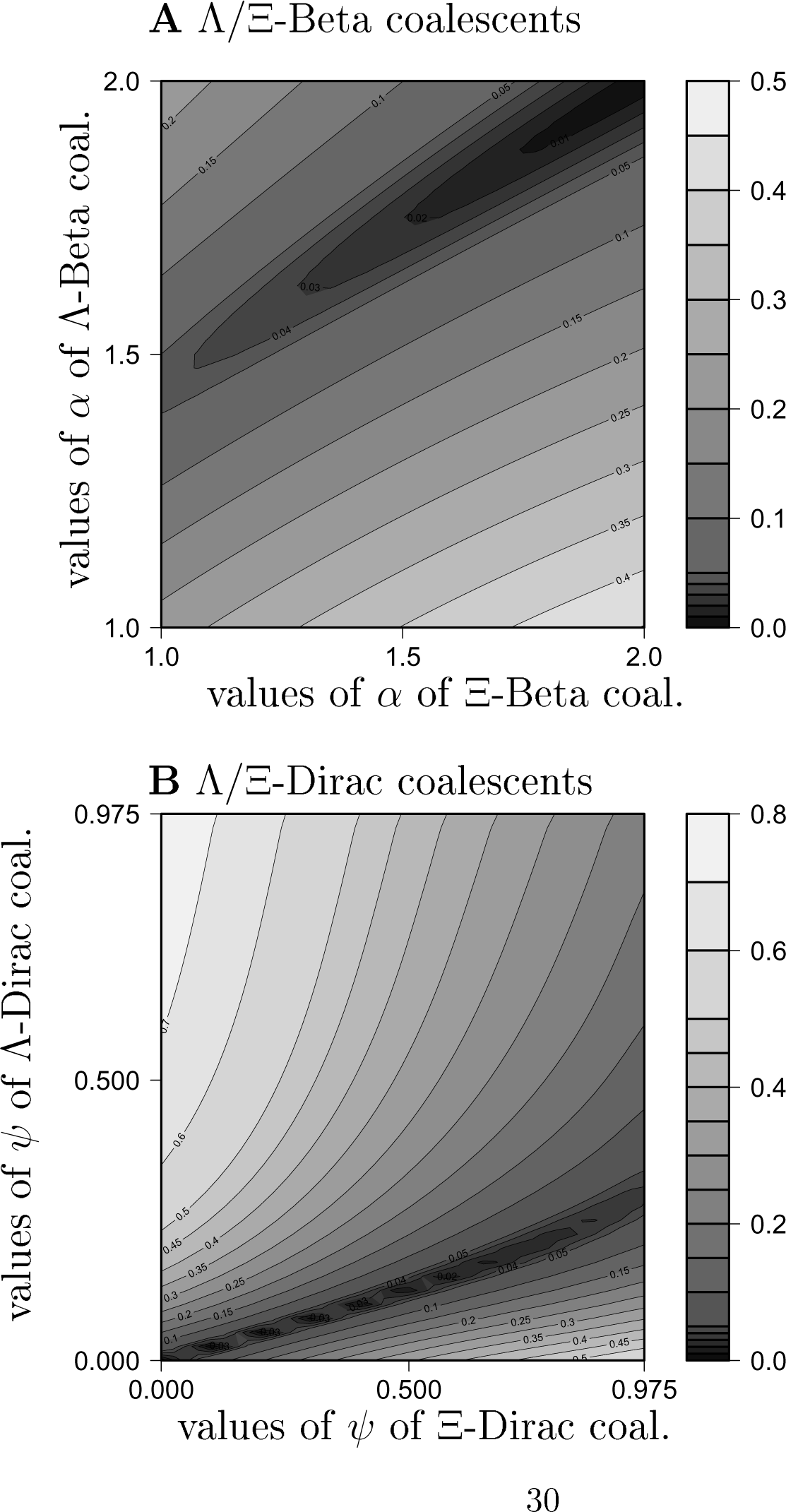
The distance between 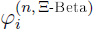 (32) and 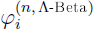 (**A**) as quantified by the ℓ_2_ norm (33); the distance between 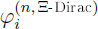 and 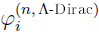 (**B**) as quantified by the ℓ_2_ norm. The number of leaves *n* = 50. Values were computed over the grid 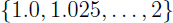 when associated with the Beta-coalescent (**A**); and 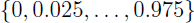 when associated with the Dirac-coalescent (**B**).

The difference in estimates between corresponding Lambda- and Xi-coalescent may be Understood from the way the Xi-coalescent process is constructed. Indeed, given *k* blocks drawn from the associated Lambda-coalescent, we see one *k*-merger with probability 4^1–*k*^, which quickly becomes small as *k* increases. A much more likely outcome is for blocks to become (evenly) distributed into four groups. The effect on the genealogy of drawing a large number *k* is thus reduced in a Xi-coalescent relative to the corresponding Lambda-coalescent.

C code written for the computations is available at http://page.math.tu-berlin.de/~eldon/programs.html.

### Application to Atlantic cod data

Atlantic cod is a diploid highly fecund marine organism, whose reproduction is potentially characterised by a skewed offspring distribution [1, 2]. Since Xi-coalescents can arise from diploid population models which admit skewed offspring distributions [34, 9], one should analyse population genetic data of autosomal loci in diploid highly fecund populations with Xi-coalescent models. Indeed, [2] obtain population genetic data at three autosomal loci ? from Atlantic cod. We use the ℓ_2_-norm (33) to fit (see Table 3) the 4-fold Xi-Beta and 4-fold Xi-Dirac coalescents to the unfolded site-frequency spectrum (USFS) of *Ckma, Myg*, and *HbA2* genes obtained by [2]. The 4-fold Xi-coalescents correspond to a diploid population in which one successful pair of parents contributes offspring in each generation.

**Table 3:** Parameter estimates 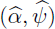 obtained by minimising the ℓ_2_-norm (33) for the unfolded site-frequency spectrum of Atlantic cod data [2]. Parameter α associated with the Xi-Beta and the Lambda-Beta coalescent was estimated over the grid 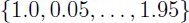, and ψ associated with the Xi-Dirac and the Lambda-Dirae coalescent was estimated over the grid 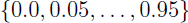.

| locus | sample size | seg. sites | Xi-coalescents |  |  |  |
| --- | --- | --- | --- | --- | --- | --- |
| | | | $\hat{\alpha}$ | $\ell_2(\hat{\alpha})$ | $\hat{\psi}$ | $\ell_2(\hat{\psi})$ |
| *Ckma* | 122 | 91 | 1.0 | 0.08 | 0.30 | 0.10 |
| *CkmaA* | 43 | 47 | 1.0 | 0.21 | 0.65 | 0.12 |
| *CkmaB* | 79 | 55 | 1.0 | 0.18 | 0.50 | 0.13 |
| *Myg* | 45 | 29 | 1.0 | 0.27 | 0.80 | 0.14 |
| *HbA2* | 114 | 11 | 1.0 | 0.34 | 0.80 | 0.13 |
| Lambda-coalescents |  |  |  |  |  |  |
| *Ckma* | 122 | 91 | 1.25 | 0.07 | 0.05 | 0.12 |
| *CkmaA* | 43 | 47 | 1.1 | 0.13 | 0.15 | 0.11 |
| *CkmaB* | 79 | 55 | 1.1 | 0.08 | 0.15 | 0.13 |
| *Myg* | 45 | 29 | 1.0 | 0.10 | 0.20 | 0.14 |
| *HbA2* | 114 | 11 | 1.0 | 0.19 | 0.25 | 0.13 |

The USFS of the autosomal genes *Ckma, Myg*, and *HbA2* are all characterised by a high relative amount of singletons. Thus, singletons have the most weight in our estimate, in particular since we do not calibrate the difference between observed and expected values by the variance. The estimates of aα associated with the Xi-Beta coalescent are therefore all at 1.0, which we attribute to the excessive amount of singletons. The excessive amount of singletons also increases the estimate of ψ associated with the Xi-Dirac coalescent. In particular, our Xi-based estimates of ψ are higher than the Lambda-based estimates (Table 3). Possibly the Xi-coalescent assigns less mass to the external branches than the corresponding Lambda-coalescent for a given parameter value, but the exact shift in mass may vary between different Xi-coalescents. The Xi-Dirac coalescent is able to predict the excessive amount of singletons, the Xi-Beta coalescent much less so (Figures 5-6).

**Figure 5:**
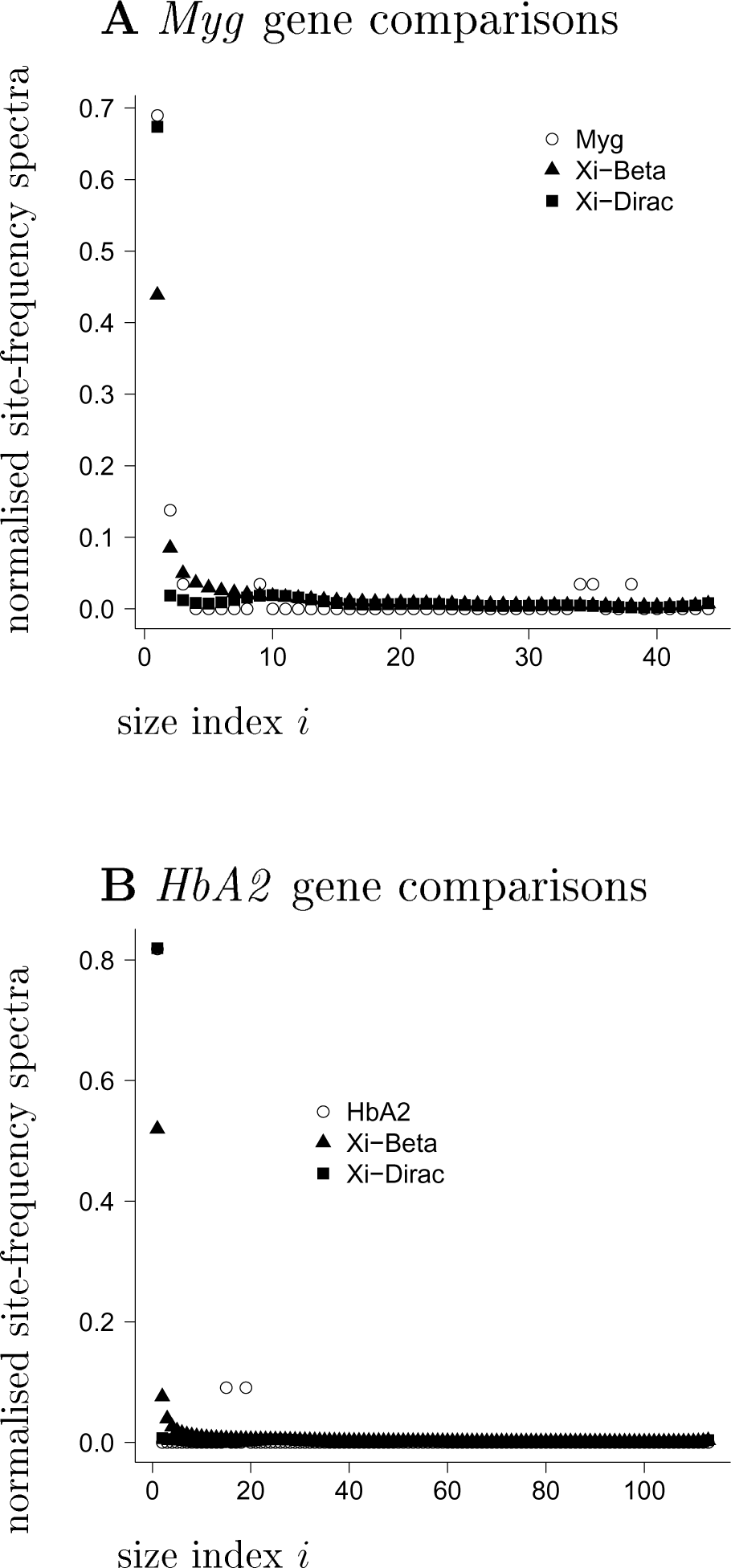
Comparison of observed 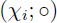 and expected (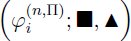) (32) normalised site-frequency spectrum for *Myg* and *HbA2* [2] with 4-fold Xi-coalescent process as shown. The associated parameter values are given in Table 3. In **A**, the observed and expected valus for the *Myg* gene are compared; in **B** for the *HbA2* gene.

**Figure 6:**
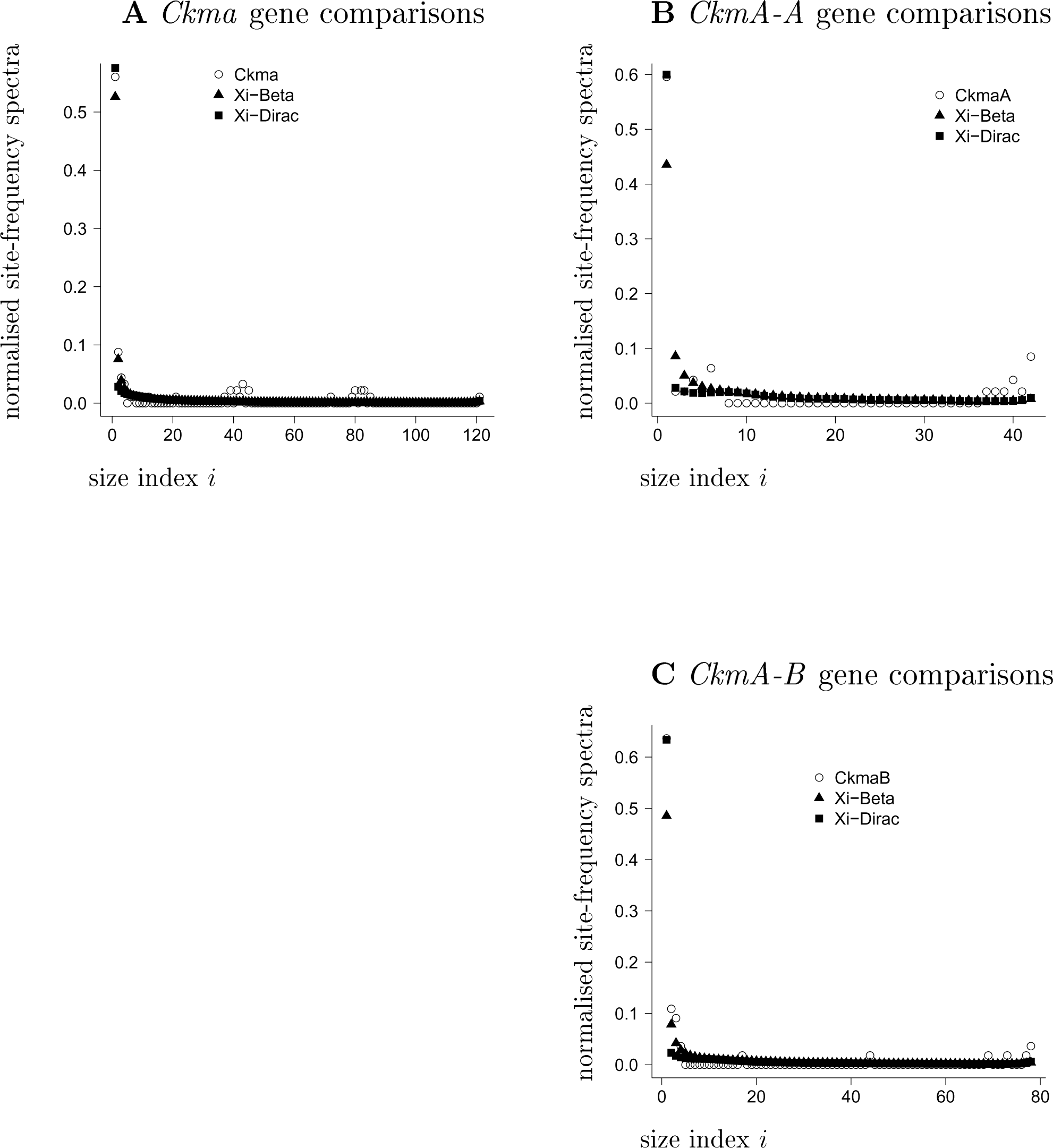
Comparison of observed 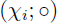 and expected (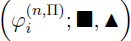) (32) normalised site-frequency spectrum for *Ckma* [2] with 4-fold Xi-coalescent process as shown. The associated parameter values are given in Table 3. In **A**, all the *CkmA* sequences are included; in **B**, only the *A* allele is considered; in **C**, only the *B* allele is considered.

Our estimates of ψ for the combined data on *Ckma* are smaller than for the partitioned data (into *A*and *B* alleles), and for the supposedly neutral loci *Myg* and *HbA2*. [2] also observe a similar pattern. Of the three loci, the Xi-coalescents give best fit to the *Ckma* data. In view of the modes in the right tail of the USFS for *Ckma*, [2] conclude that *Ckma* is under strong selection. Even though the Xi-Dirac coalescent does show multi-modal spectrum (Figure 3), the modes are small relative to the expected length of external branches, and do not quite explain the modes observed for *Ckma* (Figure 6).

## Discussion

We prove recursions for the expected site-frequency spectrum associated with Xi-coalescents which admit simultaneous multiple mergers of active ancestral lineages (blocks of the current partition). We give a class of Xi-coalescents which is ‘driven’ by a finite measure (a Lambda-measure) on the unit interval, which determines the law of the total number of active lineages which may merge each time. This class of Xi-coalescents can be applied to populations of arbitrary ploidy. We apply the recursions to compare estimates of coalescent parameters between Lambda- and Xi-coalescents. Finally, we estimate coalescent parameters associated with Xi-coalescents for Atlantic cod where the data are the unfolded site-frequency spectrum on autosomal loci.

The framework we develop will allow us to extend the recursions to more complicated scenarios, such as populations structured into discrete subpopulations [21, 31], or possibly by considering some sort of continuous distribution in space [3]. However, one would extend the recursions to such more structured frameworks at the high risk of increasing their computational complexity.

Computing the full expected site-frequency spectrum for a 4-fold Xi-coalescent with sample size *n* ≥ 100 takes unfortunately a bit of time (on the order of hours). We do not provide detailed analysis, but the time can be shortened by considering the lumped site-frequency spectrum, where one would collect all classes of size larger than some number *m* ≪ *n* into one class. How such lumping would affect the inference remains to be seen. However, one can analyse small samples (*n* ≤ 100) with our recursions, as we provide an example of. Exact likelihood methods in the spirit of [8] are yet to be developed for Xi-coalescents, and will likely be computationally intensive. In a very recent preprint, [43] introduce a different method to compute the expected SFS associated with Xi-coalescents. The method of [43] is claimed to be approximately *n*^2^ faster than our method, where *n* is sample size.

Our simple method of minimising the deviation between observed and expected values of the unfolded site-frequency spectrum should not be applied in a formal test procedure, since we do not scale the deviations by the corresponding variance. We do not give recursions for the (co)-variances of the site-frequency spectrum associated with Xi-coalescents, as these will very likely be too computationally intensive to be useful. This has already been shown to be the case for the much simpler Lambda-coalescents [10]. Our recursions provide a way to distinguish between Xi-coalescents and other demographic effects such as population growth, with the use of approximate likelihoods [22].

We obtain estimates of coalescent parameters associated with Xi-coalescents for data on autosomal loci of Atlantic cod [2]. Our estimates differ from previous estimates obtained with the use of Lambda-coalescents [2], due to the simultaneous merger characteristic of Xi-coalescents. Regardless of exact estimates, our results, coupled with those of [2], suggest that multiple merger coalescents might be the proper null model with which to analyse population genetic data on Atlantic cod, as well as other populations which may exhibit high fecundity coupled with skewed offspring distribution (HFSOD). Our specific examples of Xi-coalescents may well represent an oversimplification of the actual mating schemes; ie. assuming only one successful couple contributes offspring in each generation. However, one could use the framework of [14] to develop new examples of Xi-coalescents for specific mating schemes, for example when one successful female produces offspring with many males.

Rigorous inference methods to distinguish the effects of HFSOD from selection are yet to be developed. The common notion is that selective sweeps lead to an excess of singletons. The main genetic signature of HFSOD is also an excess of singletons. The unfolded site-frequency spectrum of the *Ckma* gene [2] is trimodal, with an excess of singletons, and small modes of mutations of larger size. These smaller modes are not captured by the examples of Xi-coalescents that we apply. Durrett and Schweinsberg [42, 20] use a stick-breaking construction to obtain a good approximation 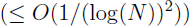 to a selective sweep where *N* denotes the population size. [2] conclude that *Ckma* is under a form of balancing selection. Our examples of Xi-coalescents also give best fit to the *Ckma* gene of the three loci studied by [2], although our method does not constitute a formal test. We refer to [2] for more detailed discussion of the variation observed at *Ckma*, and the supposedly neutral nuclear loci *Myg* and *HbA2*. The congruence between our Xi-coalescent examples and *Ckma*, and between Xi-coalescents and selective sweeps studied by [42, 20], and the different site-frequency spectra predicted by Lambda- and Xi-coalescents, leads us to conclude that Xi-coalescents form an important class of mathematical objects.

## ACKNOWLEDGMENTS

JB and BE acknowledge support by Deutsche Forschungsgemeinschaft (DFG) grant BL 1105/3-1 as part of SPP Priority Programme 1590 ‘Probabilistic Structures in Evolution’.

## Appendix

### Proof of recursion (26) for 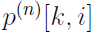 (Thm. 2)

The requirement 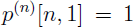 is obvious (therefore 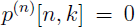 for all *k* > 1). From now on, we consider the case 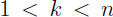. By 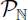 we denote the set of all partitions of 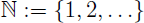. Let 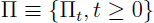 be a 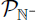-valued exchangeable coalescent, defined on the probability space 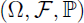. By 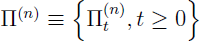 we denote the projection of Π onto 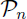, which will be associated with coalescent processes started from *n* ≥ 2 leaves. Define,

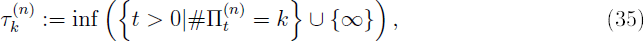

ie. the first time the block counting process associated with Π(*^n^*) hits state *k*, with 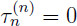.

Let

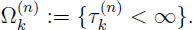

Recall that we assume that 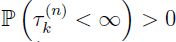 for each 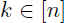. Thus we can define the conditional law 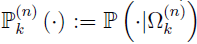. On the conditional probability space 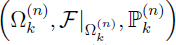, we sample uniformly at random a block π_0_ from the blocks of 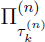, ie. 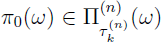 for 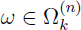. Then we can write

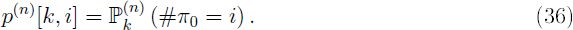

We define the first jump time τ of the block-counting process

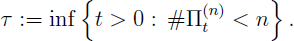

Now consider some 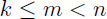, and partition 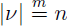, ie. an integer partition ν of *n* into *m* elements. Define

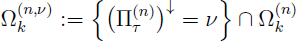

(recall that 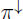 denotes the integer partition associated to 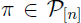 obtained by listing the block sizes of π in decreasing order). Then it is clear that we have the decomposition

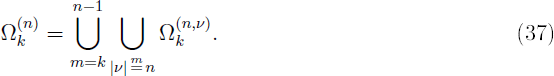

We define the conditional law 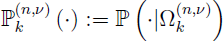.

For each 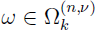, we define the set of blocks

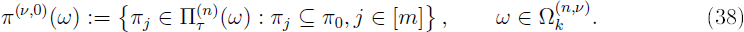

The set of blocks 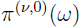 contains all blocks of the partition 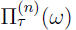 that will eventually merge into the block 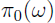

For any integer subpartition 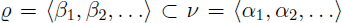, we need to be able to compute the probability 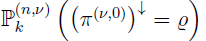 on 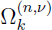. Indeed,

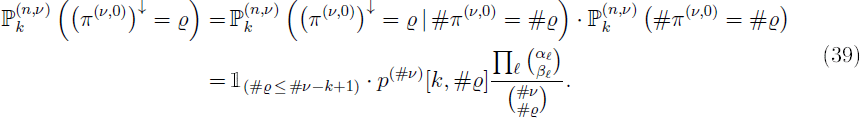

The event 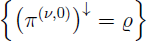 states that the sizes of the blocks of 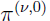 are given by the integer subpartition ϱ. Equation (39) follows from the exchangeability of П, and the form of the probability density function of the multivariate hypergeometric distribution, which applies to the event of sampling βℓ blocks from αℓ for each ℓ for a total of #ϱ out of #ν The condition 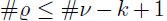 is required since we condition on the block counting process *k* blocks.

Given (39) we can compute 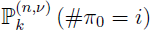. Indeed, the decomposition

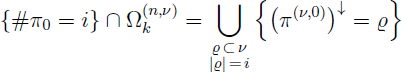

(where the union is disjoint) together with (39) gives

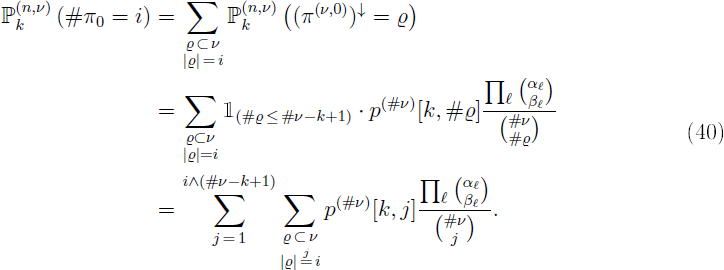

Given Equation (40) one can now compute 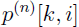: by the decomposition (37), we obtain

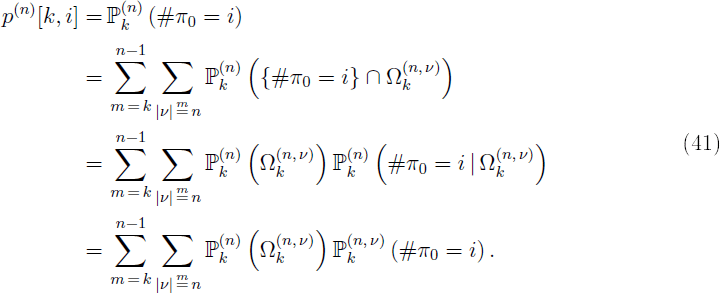

We apply Lemma 5 (stated and proved below) to 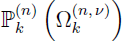 and Equation (40) to 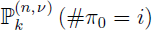 to obtain

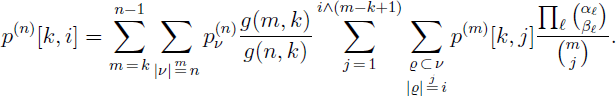

### Proof of computation of 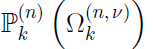 (Lemma 5)

Let 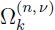, 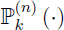, 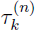 (35) and τ be as defined in the proof for Thm. 2. Recall further that 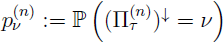, see Equation (25).

By 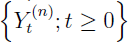, 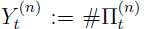 we denote the block-counting process associated with Π^(*n*)^, starting from *n*. Let *g*(*n, k*) denote the expected length of time that 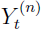 spends in state *k* ≤ *n*,

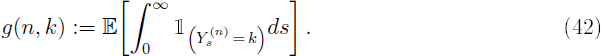

Clearly, 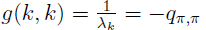 where 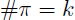, see (12).

**Lemma 5.** *For* 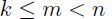 *and* 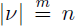, *assuming* 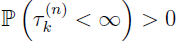, *we have*

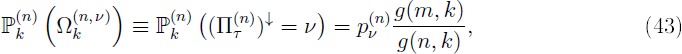

*where g (.,.) is defined in* (42).

**Proof.** Assuming 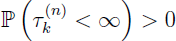, one obtains

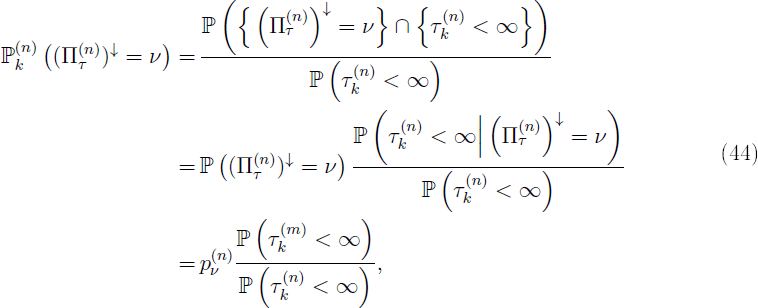

where we use #ν = *m* and apply the strong Markov property of the process 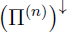 at time τ to obtain the last equality.

Now we consider 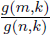, and obtain

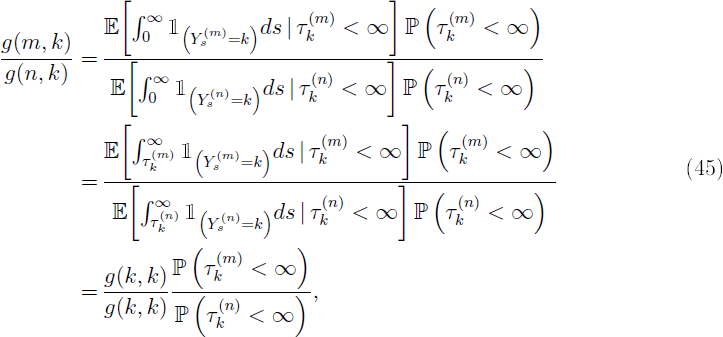

where we use the strong Markov property of the block-counting process in the last step. Now the statement follows from (44) and (45).

### Proof of recursion (23) for *g(n,k)* (Lemma 1)

**Proof.** For *n* ≥ 2, let 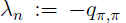 with *q*_ππ_ given by (12) and #π = *n*. Again write 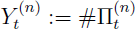 for the block-counting process starting from *n*.

Let 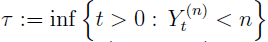 denote the first jump time of the block-counting process, and let 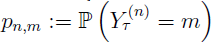, 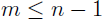. For *k = n*, one obtains

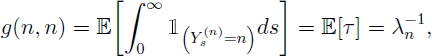

and (24) is established. For *k < n*, we decompose according to the value of the block-counting process after the first jump and obtain

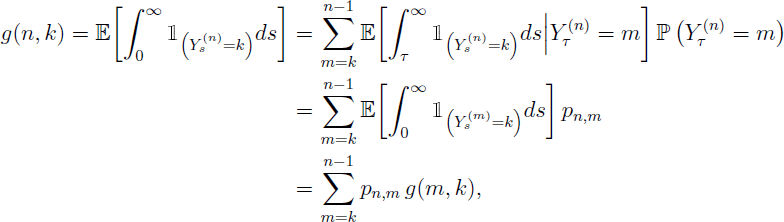

where we use the strong Markov property of *Y* in the second-to-last equality. Thus (23) is established.

### Proof of Equation (14)

Fix *m* ≥ 2, 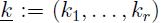 with 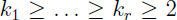, 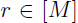 and let 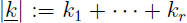, 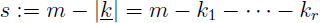. Then we have to prove that

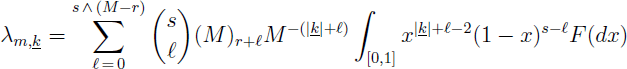

Let Δ denote the infinite simplex, and recall the function *f* from (7) used to describe the rates (12) of a Xi-coalescent. We define the map 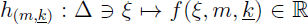. We then have that the rate of a 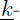merger is given by

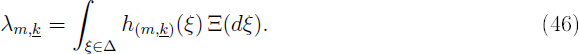

Since supp 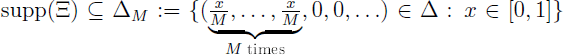, we can rewrite the rate in (46) as

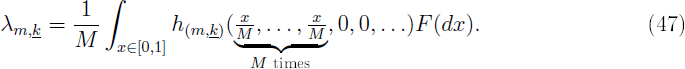

Define 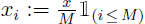. Then the integrand in (47) is given by

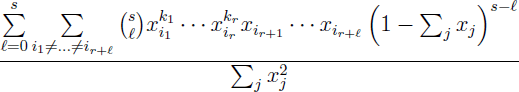

which is zero if 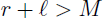. For 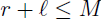, one obtains

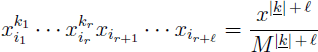

for any choice of indices 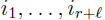 which are all different and smaller or equal to *M*. Therefore, with 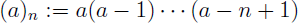 for 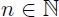, 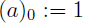 denoting the falling factorial, we have

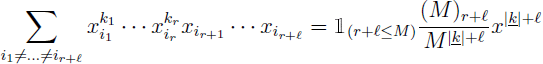

We note further that

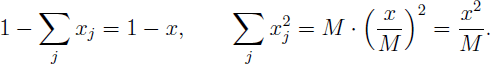

Thus we get

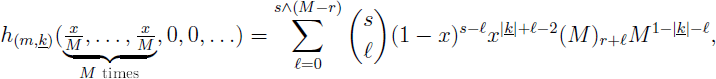

and we can represent the rate 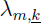 as

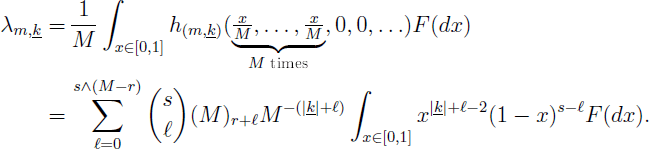

This completes our proof.

### A condition for the block-counting process to hit state *k* (Propositon 6)

We now give a necessary and sufficient condition for the block-counting process of a Ξ-*n*-coalescent to hit state 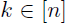 with positive probabilty, i.e. for 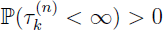, where 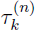 is defined as in (35).

**PROPOSITION 6.** *Let n, k be positive integers satisfying 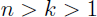, and let 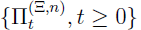 be a Ξ-n-coalescent. Let Δ denote the infinite simplex* (6), *and 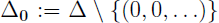. Then 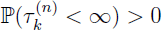 holds if and only if* 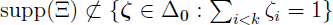.

Coalescents that fail to satisfy the condition 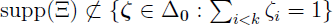 must have the property that the *first* merger-event will almost surely result in a state of less than *k* blocks, regardless of the value of *n*. Hence it FOLLOWS that any *m*-coalescent which has a ‘Kingman part’ (i.e. Ξ has mass at (0, 0,…)), or which admits at least *k* simultaneous mergers, must necessarily satisfy 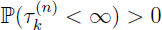.

**Proof.** We consider the decomposition 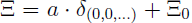 into a Kingman part and a non-Kingman part, as used earlier in the paper.

In the case *a* > 0, the condition 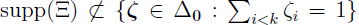 is satisfied since 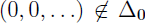. Since there is a non-zero probability of the first *m* – *k* merger-events all being single binary mergers, it follows that 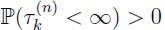 must hold in this case.

In the remainder of this proof, we shall concern ourselves with the case *a* = 0. We begin by considering the Poisson process construction of the Ξ-coalescent as outlined in [11, section 1.4]. This correspons to letting 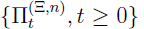 be driven by a Poisson point process *𝒩* on 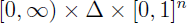 with intensity-measure

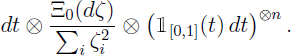

From a realization of the process *N*, we then construct 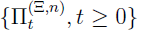 by considering the atoms of 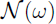 in in ascending order of the first coordinate. Given 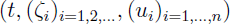 and 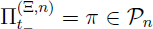, we construct 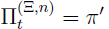 as follows:

- For 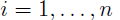 let 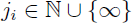 be defined by

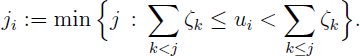
- For *J* = 1, 2, …(∞ *not* included), let

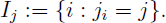

Let π*_i_* denote the *i*th block of π in order of least elements, and for all *j* = 1, 2, … merge all blocks with an index belonging to *I_j_*, in order to obtain π'.

In the following, we let

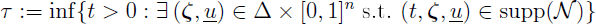

be an *N*-measurable stopping time corresponding to the time until the first event of *N*.^1^ Since Ξ is a nont-trivial measure, it follows that τ < ∞ holds almost surely. furthermore, it follows by construction that 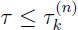.

We first consider the case 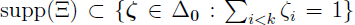. It follows immediately from the Poisson point process construction that 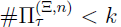 must hold (and by extension 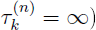, since every block of 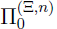 must merge into one out of strictly less than *k* groups of blocks at time τ. This proves the ‘only if’-part in Proposition 6.

In order to show the ‘if’-part in Propostion 6, suppose now that 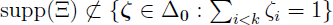. We consider the two 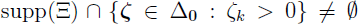 and 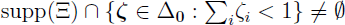.

In the case 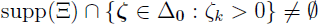, let 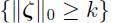 be shorthand for the event

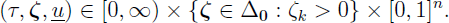

We now aim at giving a non-trivial lower bound on 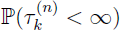 by bounding the right side of the following inequality from below:

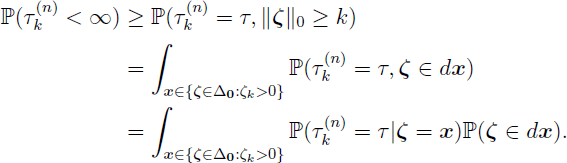

The inequality follows since the event 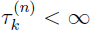 is a consequence of the event 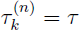, and τ < ∞ holds almost surely. The equalities on the other hand are an immediate consequence of the poisson point process construction. Hence we may now devote ourselves to proving that the last integral is positive:

Let 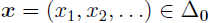, and let *x_k_* > 0. Since

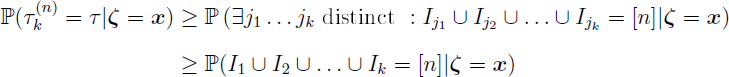

is true by construction, and

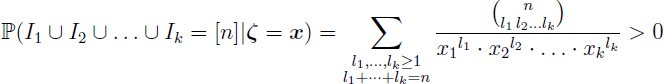

holds, it follows that

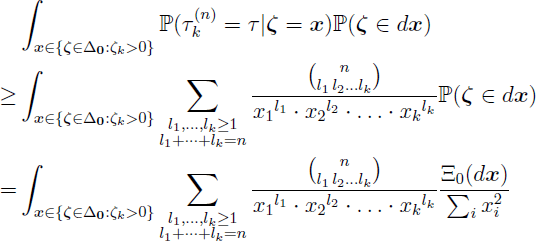

must also hold. Since the map

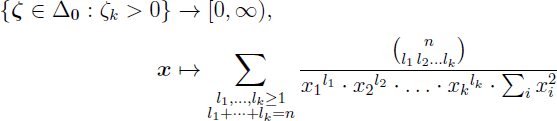

is strictly positive and 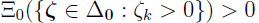 holds by assumption, it follows that

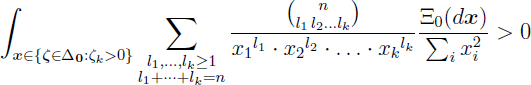

must hold. This in turn establishes a non-trivial lower bound on 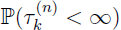.

In the case supp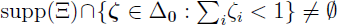, we again rely on the Poisson point process construction. For 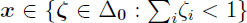, denote 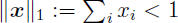. Let 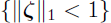 be shorthand for the event

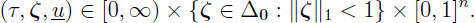

In a manner analogous to the previous case, we can establish that

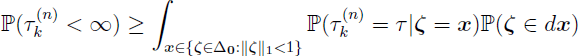

holds. Again, the integrand is strictly positive. This time it is bounded from below by

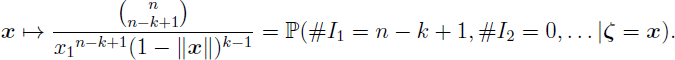

Since 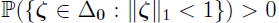 holds by assumption, this again establishes a non-trivial lower bound on the right side of the above inequality.

### Verification of the ℓ_2_ norm (33)

**Table 4:**
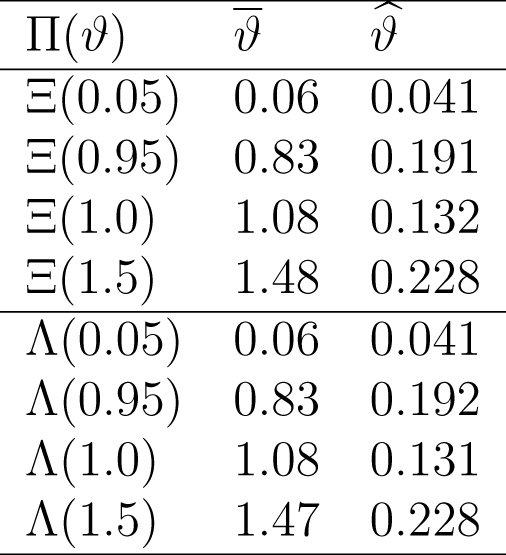
Verficiation of the ℓ_2_ (33) for estimation of coalescent parameters ν associated with Λ or Ξ-coalescents (Table 2). The ‘data’ are branch lengths associated with given coalescent, and parameter estim ates (mean 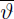; standard deviation 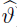) are given for the same coalescent using the ℓ norm. Based on number of leaves *n* = 50, and 10^5^ replicates.

## Footnotes

1 The definition of τ is in terms of *N*, and *not* in terms of 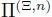. As a consequence, we may have 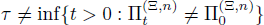.

